# ESM-Effect: An Effective and Efficient Fine-Tuning Framework towards accurate prediction of Mutation’s Functional Effect

**DOI:** 10.1101/2025.02.03.635741

**Authors:** Moritz Glaser, Johannes Brägelmann

## Abstract

Predicting functional properties of mutations like the change in enzyme activity remains challenging and is not well captured by traditional pathogenicity prediction. Yet such functional predictions are crucial in areas like targeted cancer therapy where some drugs may only be administered if a mutation causes an increase in enzyme activity. Current approaches either leverage static Protein-Language Model (PLM) embeddings or complex multi-modal features (e.g., static PLM embeddings, structure, and evolutionary data) and either (1) fall short in accuracy or (2) involve complex data processing and pre-training. Standardized datasets and metrics for robust benchmarking would benefit model development but do not yet exist for functional effect prediction.

To address these challenges we develop ESM-Effect, an optimized PLM-based functional effect prediction framework through extensive ablation studies. ESM-Effect fine-tunes ESM2 PLM with an inductive bias regression head to achieve state-of-the-art performance. It surpasses the multi-modal state-of-the-art method PreMode, indicating redundancy of structural and evolutionary features, while training 6.7-times faster.

In addition, we develop a benchmarking framework with robust test datasets and strategies, and propose a novel metric for prediction accuracy termed relative Bin-Mean Error (rBME): rBME emphasizes prediction accuracy in challenging, non-clustered, and rare gain-of-function regions and correlates more intuitively with model performance than commonly used Spearman’s rho. Finally, we demonstrate partial generalization of ESM-Effect to unseen mutational regions within the same protein, illustrating its potential in precision medicine applications. Extending this generalization across different proteins remains a promising direction for future research. ESM-Effect is available at: https://github.com/moritzgls/ESM-Effect.

## 1 Introduction

Current approaches to pathogenicity prediction of mutations focus on classifying mutations as either benign (without effects on the protein) or pathogenic/damaging. However, they fail to quantify functional effects (e.g., changes in activity) or distinguish activating from inactivating mutations. Yet he exact functional effect is crucial for protein engineering and precision medicine where specific cancer drugs can only be administered when a mutation causes an increase in enzyme activity (Leichsenring et al., 2019; Mateo et al., 2022). This challenge is further exacerbated by the rapid increase in mutations identified in routine patient sequencing, driven by the decreasing cost of sequencing technologies (Pasmans et al., 2021). While Deep-Mutational Scans (DMS, i.e., experimentally testing effects of all possible mutations in a given protein) offer clinicians precise functional insights, they are laborious, costly and often fail to cover the full protein of interest (Karczewski et al., 2020). These limitations underscore the need for accurate computational methods to efficiently predict the functional effect of mutations.

Pathogenicity predictors struggle to accurately predict the multi-faceted functional effects of specific mutations, such as rare gain-of-function enzyme mutations (Appendix 7.1), because they adopt a generalist strategy, scoring all possible variants across the (human) proteome (Cheng et al., 2023). This limitation arises from the biological complexity and specificity required for such tasks, which cannot be reliably captured by large-scale pre-training and the current architectures (Livesey & Marsh, 2023). Thus, functional effect predictors have been developed which are fine-tuned on DMS datasets achieving higher protein-specificity and accuracy. Yet they are naturally limited to unseen mutations and regions in their training protein which is still valuable to some application settings. Current approaches such as Rep2Mut-V2 and PreMode rely on non-fine-tuned embeddings or intricate multi-modal features requiring complex pre-processing and pre-training (Derbel et al., 2023; Zhong et al., 2024).

In this paper, we address these limitations by

1. first evaluating the shortcomings and potential of existing methods for both pathogenicity and functional effect prediction and
2. then developing the optimal framework for ESM2-based functional effect prediction through detailed ablations of various fine-tuning strategies and prediction head architectures to identify the most performant ones. Based on these insights, we propose the ESM-Effect framework, which achieves state-of-the-art (SOTA) performance on functional effect predictions slightly outperforming multi-modal competitors at 6.7x faster training speed.
3. Finally, we analyze the strengths and weaknesses of ESM-Effect’s generalization capabilities and propose robust benchmarks to facilitate further progress in the field.

## 2 Background

### Mutation Effect Prediction as a Question of Pathogenicity

Mutations affect proteins by diverse means, rendering precise measurement of their impact challenging. To simplify, the concept of “mutation pathogenicity” categorizes mutations as either “pathogenic” (i.e., disrupting physiological protein function) or “benign.” Pathogenic mutations can reduce organism fitness and are rare in natural sequences, such as those in UniRef (Suzek et al., 2007), representing the physiological sequence space. Models learn pathogenicity from large datasets of natural sequences, scoring the likelihood of mutations based on their presence in (physiological) evolutionary or multiple-sequence alignments (MSAs) (Meier et al., 2021). However, this broad definition oversimplifies the heterogeneous functional effects mutations can exert. For example, pathogenic mutations in an ion channel might either increase or decrease affinity (Kullmann & Hanna, 2002), whereas pathogenic mutations in kinases can increase (Gain of Function, GoF) or decrease (Loss of Function, LoF) their catalytic activity (Iyer et al., 2023).

### Mutation Effect Prediction as a Question of Functional Effects

In contrast, functional effect prediction considers a wider range of impacts, such as catalytic activity, which are more directly applicable to precision medicine and protein engineering. However, achieving high accuracy requires both protein-specific supervised data (Zhong et al., 2024) and appropriate architectures (including training strategies).

## 3 Related Work

### 3.1 Models for functional effect prediction

Existing models extend pathogenicity predictors (Appendix 7.1): Derbel et al. (2023) and Marquet et al. (2022) use static embeddings from the ESM family of PLMs (Rives et al., 2021; Lin et al., 2023) combined with a neural network head to predict functional effects from DMSs. Saadat & Fellay (2024) fine-tune ESM2 for residue-level protein sequence annotation (e.g., identifying functional features like active sites) and then classify mutations based on the probability difference of annotated features between reference and mutant sequences. These annotations are then evaluated with ClinVar data (Landrum et al., 2018). LoGoFunc, another method, performs three-class classification using a diverse feature set to make genome-wide predictions (Stein et al., 2023).

Studying the extent of the expected benefit from fine-tuning PLMs, Schmirler et al. (2024) showed that ESM2 fine-tuned with Low-Rank-Adaptation (or full fine-tuning) and a neural network regressor on top of the mean mutant sequence embeddings outperforms the simple, non-PLM baselines Homology-Based Inference and the statistical model Reference Free Analysis (RFA) on three DMS (AAV, GFP and GB1).

The latest and most complex model for functional effect prediction is PreMode (Zhong et al., 2024; Zhong & Shen, 2022), which is pre-trained on 4.7M pathogenicity-labeled mutations and then finetuned on a specific DMS. PreMode uses the static wildtype embeddings (650M ESM2 model), MSAs and additional mutation-specific features as node vectors as well as the AlphaFold2-predicted structure (AF2) for a star graph attention model (Jumper et al., 2021). PreMode outperforms a Random Forest model, pre-trained 650M ESM2 embeddings with a regression head and other adaptations of state-of-the-art pathogenicity predictors given the same input features as PreMode. Besides, the authors’ preliminary analyses showed that LoF, GoF and neutral mutations have distinct but overlapping (i.e., no unique intervals exclusive to any one class) distributions for pLDDT scores, conservation levels, and solvent accessibility.

Finally, some pathogenicity predictors can also be used for functional effect prediction, but their accuracy is limited due to their generalist nature (Jagota et al., 2023; Lafita et al., 2024).

### 3.2 Databases and existing Benchmarks for Mutation Effect Prediction

To compare predictors, large databases of clinically annotated mutations and Deep Mutational Scans (DMS) have been developed as well as numerous experimental efforts exploring and testing mutations experimentally (Backman et al., 2021; Dunham & Beltrao, 2021; Esposito et al., 2019; Exome Aggregation Consortium et al., 2016; Gao et al., 2023; The UniProt Consortium et al., 2023). Widely-used resources include the ProteinGym, which serves both as a repository for Deep Mutational Scans (DMS) and as a benchmarking platform for evaluating pathogenicity predictors (Notin et al., 2023). Similarly, MaveDB provides a curated repository of DMSs, while ClinVar includes clinical annotations with benign and pathogenic labels (Landrum et al., 2018; Rubin et al., 2021).

Livesey & Marsh (2023) used 26 DMS to benchmark 55 pathogenicity predictors reporting respectable performance (measured by Spearman correlation and AUROC) in distinguishing pathogenic variants. However, their findings underscore substantial variability across predictors, with particularly poor performance on DMSs that included GoF mutations.

## 4 ESM-Effect

### 4.1 Problem Statement

Existing methods do not fine-tune the PLM to obtain embeddings or use varying, mostly mean-based regression heads. We start the development of ESM-Effect by assessing the impact of PLM model size on performance to select the optimal base model size trading-off compute efficiency and performance. Then we perform detailed ablations of combinations of different training regimen and regression heads. Finally, we unite the most performant techniques regarding the training strategy and the regression head in ESM-Effect. In the results section we compare ESM-Effect to the multimodal SOTA PreMode which uses static embeddings, AF2 structure and MSAs to assess the benefit of its multi-modality.

### 4.2 ESM-Effect: Developing the optimal prediction architecture

#### ESM2 Model Size

Scaling laws in natural language processing (NLP) suggest that larger models are more compute-efficient for modest-sized datasets (Kaplan et al., 2020). These principles also hold in biological applications (Cheng et al., 2024), with increasing ESM2 model size leading to lower language modeling loss and better performance in structure prediction (Lin et al., 2023). To investigate whether these trends extend to the downstream task of functional effect prediction, we compared the performance of different ESM2 model sizes (8M to 3B) on AAV, GB1, and GFP DMS datasets (models trained by Schmirler et al. (2024)) with the validation perplexity from pre-training reported by Lin et al. (2023) (cf. Figure 1). We find that scaling laws do not hold in the DMS fine-tuning context. No obvious performance improvements emerge with larger models across all DMS uniformly. Consequently, we select the 35M ESM2 model due to its favorable balance of computational efficiency and performance.

**Figure 1:**
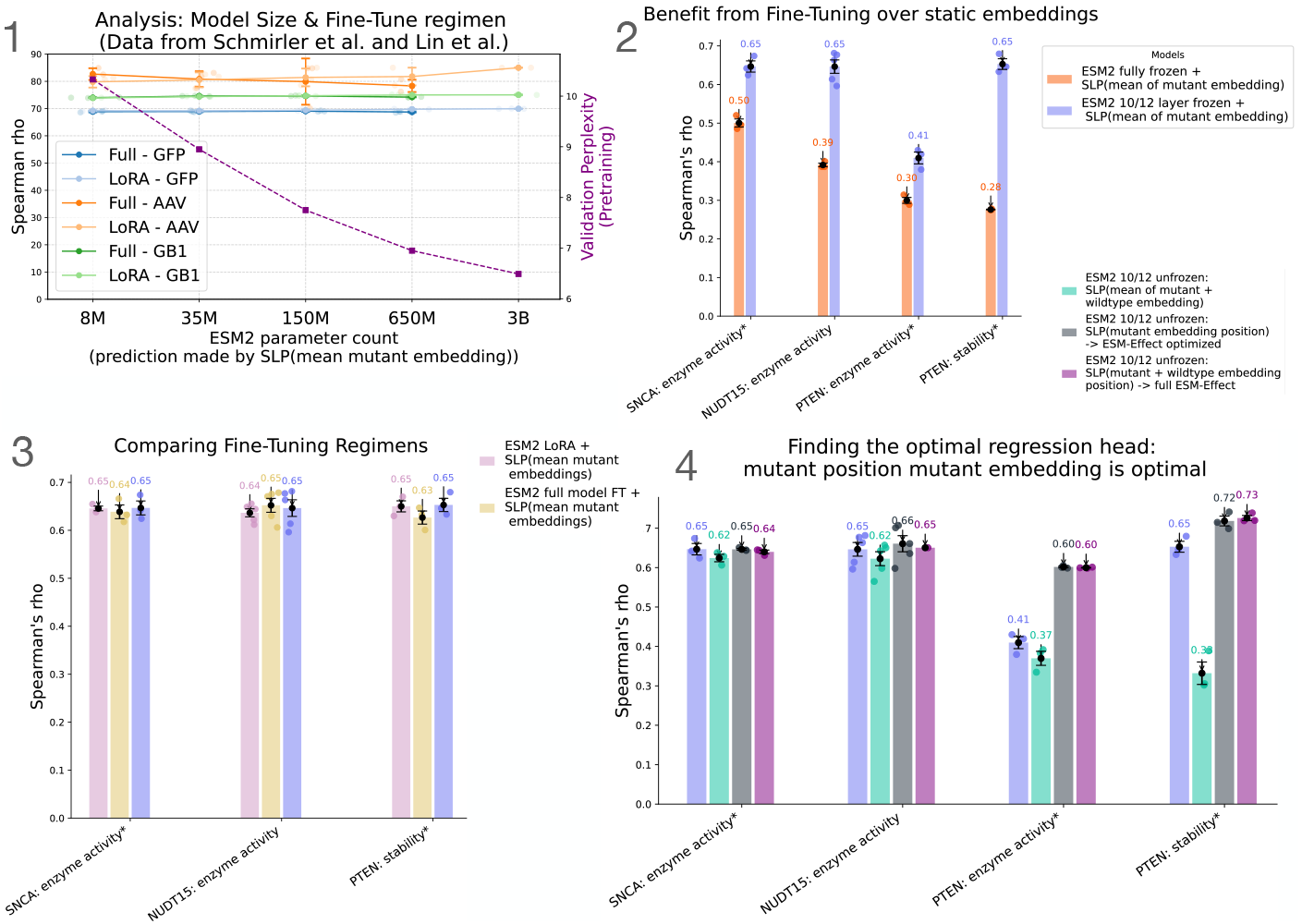
1 - Kaplan et al. (2020) scaling laws do not extend to downstream functional effect fine-tuning performance but are consistent with pre-training metrics (e.g., validation perplexity, CASP14 performance). Performance difference between fine-tuning regimens (Low-Rank Adaptation (LoRA) vs. full Fine-Tuning) is minimal. 2 - Significant benefit from fine-tuned embeddings. * indicates that only three of five available seeds were used due to resource limitations. 3 - Minimal performance differences between fine-tuning strategies. Unfreezing the last two layers was selected for ESM-Effect due to interpretability advantages etc. Information on training characteristics for the PTEN DMS is in the Appendix 7.8. 4 - Analysis of the optimal regression head. Note that mutation position based heads require a maximum of 10 epochs for optimal performance while mean based heads take longer and suffer from instable training for PTEN DMS (cf. Appendix 7.8)

#### The Value of Fine-Tuned Embeddings

Previous approaches to functional effect prediction have relied on static embeddings from fully frozen ESM models combined with various prediction heads (Marquet et al., 2022; Derbel et al., 2023; Zhong et al., 2024). To evaluate whether this limitation constrains performance, we compare static 35M ESM2 embeddings to fine-tuned 35M ESM2 embeddings (with the last two layers unfrozen) across four DMS datasets. Both approaches use a prediction head that inputs the mean of the mutant embeddings into a Single-Layer Perceptron (SLP) for a fair comparison. As shown in Figure 1, fine-tuned embeddings consistently outperform static embeddings despite dataset-specific variations. This demonstrates a critical shortcoming of existing methods and establishes fine-tuning as a key design choice for ESM-Effect.

#### LoRA, Full and Partial Fine-Tuning

Our previous analysis of the data from Schmirler et al. (2024) also confirmed that LoRA and full fine-tuning achieve comparable performance as reported by Schreiber (2023) and Schmirler et al. (2024). To independently validate this and extend the analysis, we evaluated LoRA, full fine-tuning and partial fine-tuning (unfreezing the last two layers) on three diverse DMS datasets. As shown in Figure 1, all three strategies performed equivalently.

This result diverges from findings in NLP tasks, where LoRA has been shown to underperform full fine-tuning in domains like programming and mathematics (Biderman et al., 2024). Accordingly, the functional effect prediction task exhibits unique characteristics, making LoRA and layer-freezing viable alternatives for parameter-efficient fine-tuning within the ESM-Effect framework. For further development, we selected the strategy of unfreezing the last two layers for ESM-Effect, as it achieves comparable performance while avoiding the extensive hyperparameter tuning required by LoRA (cf. Appendix 7.3). This approach offers a more straightforward implementation path while maintaining interpretability. To deeper investigate the fine-tuning behavior of ESM2 we evaluated the amount of unfrozen layers (compared to full, 10/12 frozen layers and no fine-tuning above) and the position of one unfrozen layer in the model but none influenced model performance (cf. Appendix 7.3.

#### Regression Head

Previous methods have either used the mean embedding of the mutant sequence or combined static embeddings of the mutant and wildtype sequences at the mutation position as inputs. Building on fine-tuning the 35M ESM2 model (with 10 of 12 layers frozen), we evaluated four regression head designs across four DMS datasets: (1) The mean embedding of the mutant sequence, (2) a linear combination of the mean embeddings of mutant and wildtype sequences, (3) the embedding at the mutation position of the mutant sequence and (4) a linear combination of the mutation position embeddings of mutant and wildtype sequences.

This analysis allowed us to assess the relative importance of (1) the mutation position and (2) the specific wildtype residue as landmarks of the physiological sequence space. Figure 1 shows that, while all four regression heads performed similarly for SNCA and NUDT15 DMS datasets, the mutation position-based regression head significantly outperformed mean-embedding-based approaches for the PTEN stability and PTEN enzyme activity DMS datasets. Notably, this performance gain occurred even though the second, mean-based approach incorporated information about the mutation position and wildtype residue, showing the utility of the mutation position as a valuable inductive bias for these tasks.

**The ESM-Effect Architecture** thus comprises the 35M ESM2 model with 10 of 12 layers frozen and the mutation position regression head (cf. Figure 2). The model’s performance is driven by two key inductive biases in the regression head:

**Figure 2:**
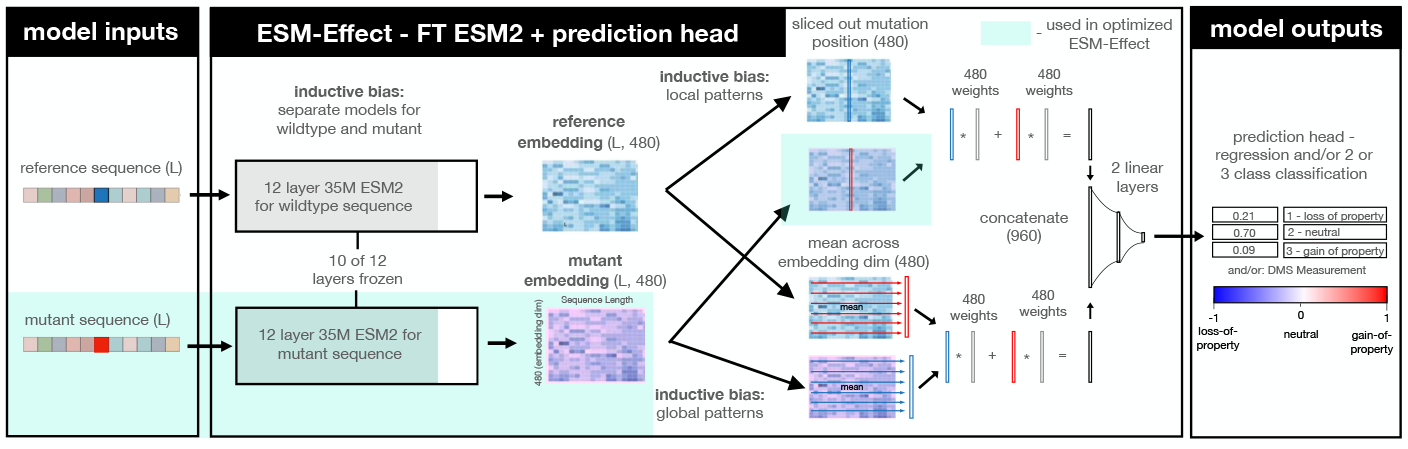
Architecture of full ESM-Effect. Embedding parts and data used for optimized ESM-Effect is highlighted in light green.

- the mutation effect is relative to a wildtype sequence
- mutation impact is largest at the mutation position

While the full architecture, incorporating both mutant and wildtype embeddings, directly implements these biases, a simpler version — using only the mutation position embedding of the mutant sequence — achieves comparable performance with approximately half the computational cost. We term this streamlined version optimized ESM-Effect. Besides, we introduce the ESM-Effect Speed implementation which collects the embeddings from the 10th layer in the first epoch, and reuses them in the remaining epochs to only forward-pass, back-propagate and train the last two layers resulting in a 8.4x speedup.

## 5 Results

### 5.1 Performance Comparison: Optimized ESM-Effect achieves SOTA Performance

To assess the performance of ESM-Effect, we compare it to the state-of-the-art method, PreMode, which is pre-trained on 147k pathogenic/benign mutation labels and fine-tuned on nine diverse DMSs. Unlike ESM-Effect, which relies solely on sequence input and its learned embeddings, PreMode incorporates static ESM2 embeddings, AF2 structures, and MSAs. In natural language processing multi-modal approaches achieve significant performance gains, and it is assumed that this extends to mutation prediction. Surprisingly, ablation analyses by removing modalities from the PreMode model revealed only marginal performance decreases when any one of the three modalities is excluded. This indicates that the information they provide for functional effect prediction is largely redundant.

Optimized 35M ESM-Effect performs at least as well as PreMode but trains 6.7x faster and does not require pathogenicity prediction pre-training (cf. Figure 3, Table 1). It also uses a fraction of PreMode’s storage (cf. Methods 7.9.2). ESM-Effect model mostly outperforms PreMode (5 of 8 DMS) by varying margins. The full ESM-Effect model and the optimized model mostly perform on par. This relates to the discussion of the arguable existence of one fixed wildtype sequence in the Appendix 7.6 and is testimony of the fact that ESM2’s own understanding of the physiological sequence space suffices and it does not require the (or “a specific”) wildtype residue. Thus, the optimized model without the wildtype residue input does not underperform the full model with this information. Moreover, we also experimented with Test-Time-Training finding mixed performance results (cf. Appendix 7.4) (Bushuiev et al., 2024).

**Table 1:** Table comparing the mean spearman rho on DMS between ESM-Effect models, PreMode and other setups on 3 or 5 seeds. Mean models in the table use the mutant sequence only.

| model<br>task name | ESM<br>Effect<br>full | ESM<br>Effect<br>optim. | ESM2 10/12<br>frozen mean | SLP<br>(embed.) | ESM2<br>LoRA<br>mean | PreMode |
| --- | --- | --- | --- | --- | --- | --- |
| ASPA: enzyme activity | **0.747** | 0.738 |  | 0.470 |  | 0.746 |
| ASPA: stability | **0.819** | 0.817 |  | 0.477 |  | 0.818 |
| CYP2C9: enzyme activity | **0.846** | 0.830 |  | 0.528 |  | 0.820 |
| GCK: enzyme activity* | **0.680** | **0.680** |  | 0.422 |  | 0.674 |
| NUDT15: enzyme activity | **0.676** | 0.661 | 0.646 | 0.491 | 0.636 | 0.658 |
| PTEN: enzyme activity* | 0.600 | **0.602** | 0.544 | 0.395 | 0.475 | 0.597 |
| PTEN: stability* | **0.726** | 0.718 | 0.653 | 0.540 | 0.650 | 0.703 |
| SNCA: enzyme activity* | 0.640 | 0.646 | **0.647** | 0.531 | 0.646 | 0.617 |

**Table 2:** Mapping of proteins to column names containing enzyme activity scores.

| Protein | column name for enzyme activity |
| --- | --- |
| SNCA | score.1 |
| CYP2C9 | score.1 |
| NUDT15 | score.2 |
| ASPA | score.2 |
| GCK | score.1 |
| PTEN | score.2 |

**Figure 3:**
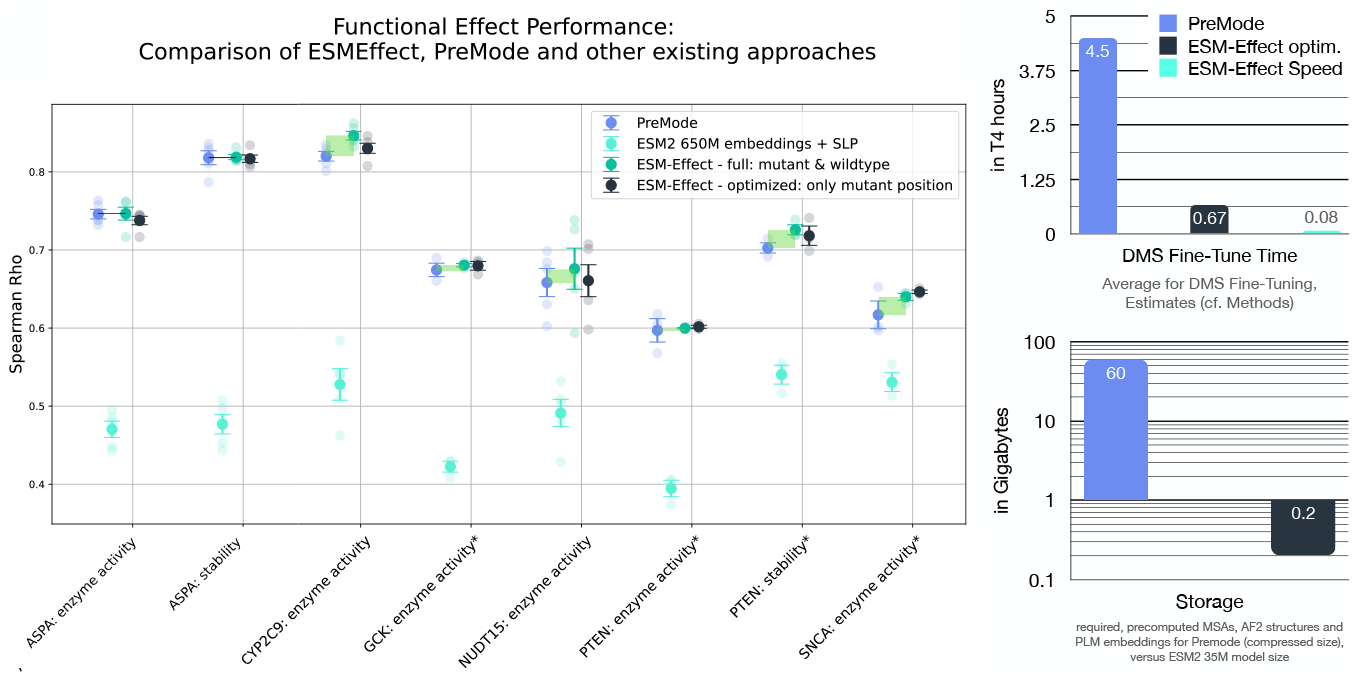
Left: Performance comparison of ESM-Effect with multi-modal PreMode. Stars indicate if 3 of 5 train-test splits were used. Right: Comparing DMS fine-tuning time and required resources of ESM-Effect (optimized version) and PreMode using estimates (cf. Methods 7.9.2).

### 5.2 Benchmarking Framework for functional effect prediction

#### General Remarks

While benchmarks are established for pathogenicity prediction, uniform benchmarks including reliable metrics and standardized testing datasets are lacking for functional effect prediction hampering useful comparisons. Thus, we propose datasets, including train-test splits, evaluation metrics, and visualization guidelines (cf. Section 5.3) to provide a more realistic framework for assessing functional effect predictors.

#### Datasets & Pre-Processing

We trained and benchmarked ESM-Effect on the same 9 DMS datasets and corresponding test splits used by PreMode, ensuring 1:1 comparability. In previous work, score calculation methods — such as normalization and aggregation of DMS experiment replicates — frequently provide insufficient information for replication. Similarly, decisions regarding the inclusion of wildtype scores and the reference sequence isoform used are not always provided. Standardizing on PreMode or miscellaneous datasets (and their respective (transformed) scores used for training) or ensuring exact sharing of datasets could address these ambiguities.

We further suggest a more rigorous testing regimen: instead of relying on random data splits, we propose evaluating models on DMS mutations from sequence intervals distinct from those in the training data. This approach provides a more realistic measure of the model’s ability to generalize to new biological contexts (see Section 5.4). For consistency, it is essential to not only share train-test splits but also the full DMS dataset and to standardize testing intervals across studies.

#### Metrics: The relative Bin-Mean Error (rBME)

For pathogenicity prediction, correlation of predicted mutation effect scores with experimental DMS scores is often evaluated using scale-invariant metrics like Spearman rank correlation, e.g., by ProteinGym. Spearman correlation is well-suited for pathogenicity because it evaluates monotonic relationships and is robust to scale differences across DMS score distributions. However, the magnitude of the Spearman’s rho may be driven by a large number of neutral mutations, while placing less weight on rare outliers. In the context of functional effect prediction this may mask biologically relevant groups of mutations with low frequency (compare Figure 5 where spearman rho for SCNA and PTEN is almost the same yet the plot reveals stark differences in performance). Functional effect prediction thus requires more nuanced evaluation that also captures rare, biologically relevant mutations.

To address this, we propose the relative Bin-Mean Error (rBME), a metric that evaluates model performance across distinct mutation effect bins, emphasizing accuracy for rare but impactful mutations (cf. Figure 4):

**Figure 4:**
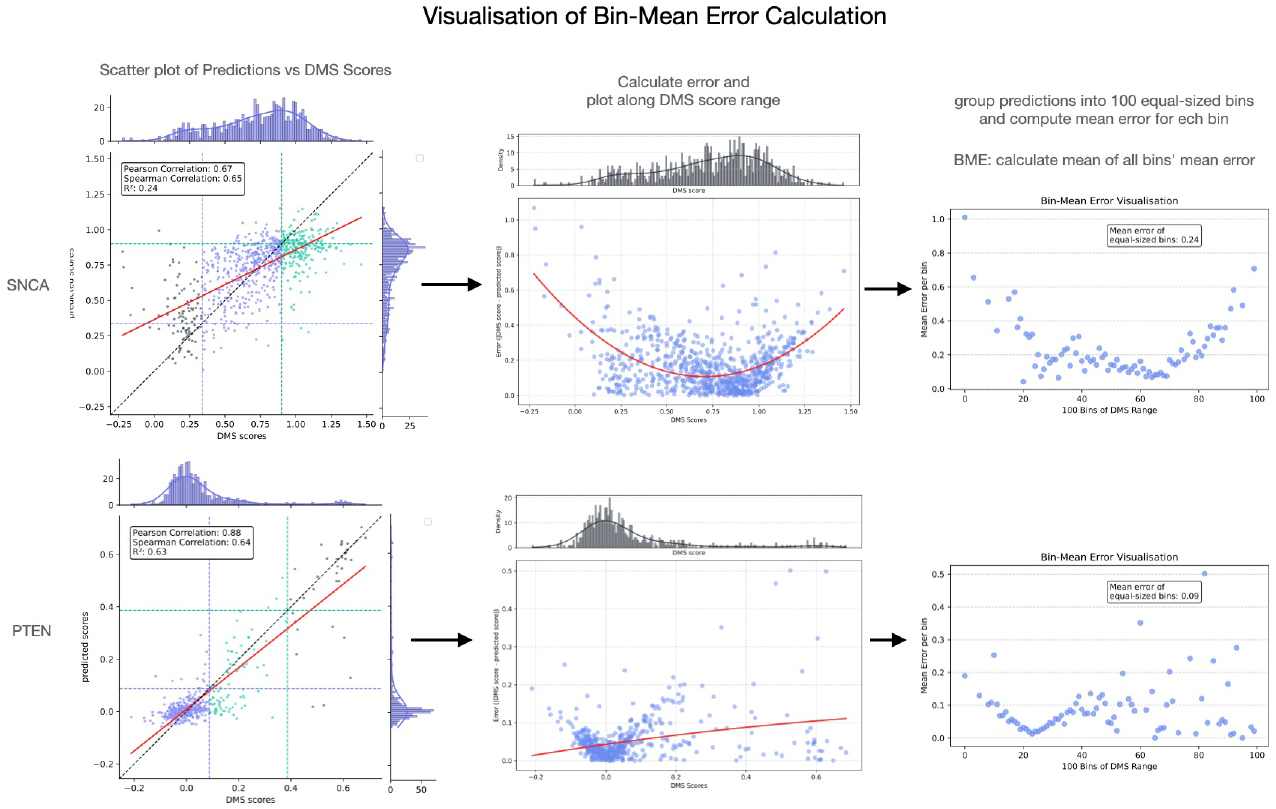
Visualization of the BME calculation steps. Predictions stem from LoRA ESM2 + SLP(mutant embeddings) fine-tuned on SNCA seed 0 and PTEN seed 1 for 20 epochs.

**Figure 5:**
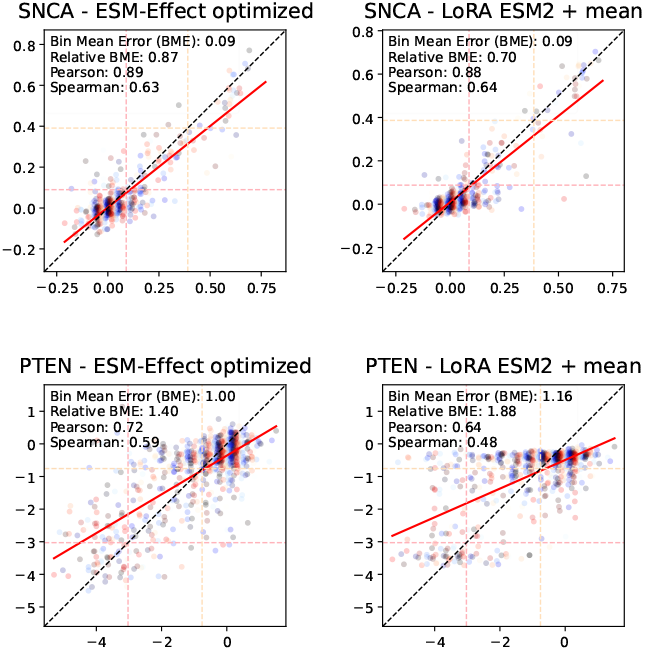
Analysis of optimized ESM-Effect and LoRA fine-tuned ESM2 with SLP(mutant mean embedding).

Let the DMS scores and predicted scores (of the test set) be denoted as *y*_*i*_ and *ŷ*_*i*_, respectively, for *i* = {1, 2, …, *N*}, where *N* is the total number of test mutations.

Define the relative error for each mutation *i* as:

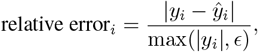

where *ϵ* is a small constant to avoid division by zero. Note that the division by *y*_*i*_ can be omitted to produce the un-normalized BME which can be used to compare different models on the same DMS. Next, group the data points into *n*_bins_ equal-width bins based on the value range of all *y*_*i*_, where *b*_*k*_ represents the *k*-th bin (typically, *n*_bins_ = 100). While the model effectively learns the true distribution of DMS scores — capturing clustered regions with many neutral mutations and producing realistic predictions — this step is crucial to mitigate metric bias and ensure balanced treatment across all regions, including easy-to-predict clusters and hard-to-predict, wider regions with rare but biologically significant Gain-of-Function mutations. The relative Bin Mean Error (rBME) is given by the mean of the mean error per bin *b*_*k*_ where |*b*_*k*_| is the number of data points in bin *b*_*k*_:

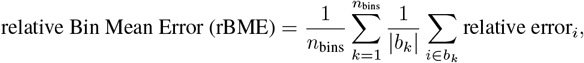

Normalization of the absolute error facilitates comparisons across different DMS, whereas the un-normalized BME metric is suitable for cross-model comparisons on the same DMS. While the optimized ESM-Effect achieves comparable Spearman correlations for PTEN and SNCA (0.59 and 0.63, respectively; cf. Figure 5), the scatter plots reveal a stark difference in performance. This discrepancy is accurately captured by the rBME metric, which reflects the disparity (0.87 vs. 1.40).

### 5.3 Prediction Analysis

While most previous studies compare prediction performance with a single metric, only plotting predictions versus ground truth truly reflects performance: a realistic plot should have the same scale for DMS scores and predicted scores axes and indicate optimal predictions with an angle bisector. Figure 5 compares the prediction characteristics of the optimized ESM-Effect model and the LoRA ESM2 model with a regression head on top of the mean mutant sequence embeddings. The prediction patterns of optimized ESM-Effect and LoRA ESM2 mean have distinct prediction characteristics, especially for PTEN enzyme activity, where it performs worse.

The prediction patterns on the SNCA DMS correlate with the high metrics (e.g., spearman rho, low BME and rBME): the models can reliably distinguish activity scores in the upper range of the DMS score distribution from scores in the lower region (score -0.2 to 0.2).

### 5.4 Investigating Generalization

As part of our proposed benchmarking framework, testing optimized ESM-Effect not by using a random split of the DMS but by using distinct sequence intervals for selecting train and testing mutations assesses generalization: the model has to infer the effect of mutations in the testing interval based on its understanding from pre-training and from patterns detected in the rest of the protein. We selected SNCA because it features a unique sequence position-to-score relationship as shown in Figure 6. Notably, the last 40 residues are predicted by MobiDB-lite to form a disordered region, lacking stable secondary structure (Necci et al., 2017).

**Figure 6:**
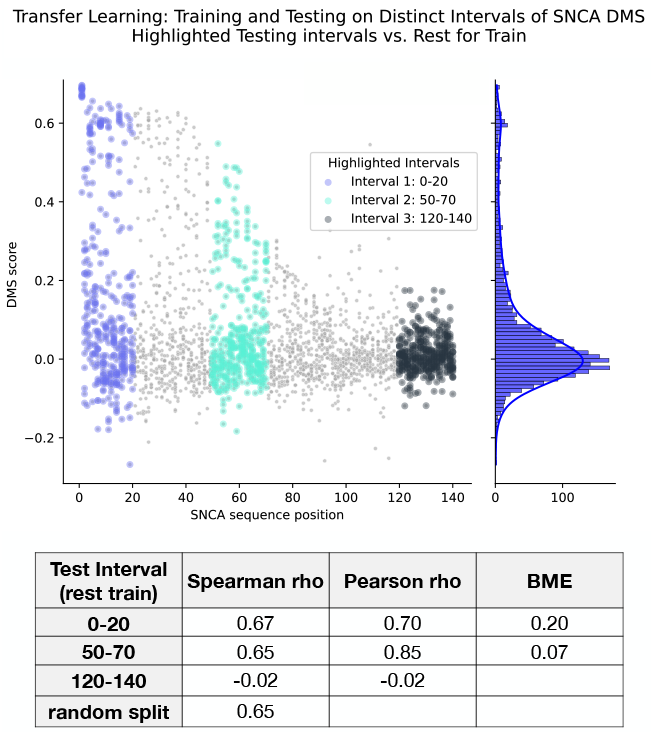
Investigating optimized ESM-Effect’s generalization capabilities on SNCA DMS. Model trained on three random seeds achieves a Spearman rho of 0.646. Each testing interval accounts for 14-15% of the total dataset, while the random split used 20%.

The generalization of ESM-Effect is highly dependent on the interval: while the model performs equally well on intervals enriched with rare, high-score mutations compared to random splits (spearman rho 0.65 vs. 0.65), it struggles within the disordered interval without these mutations (Spearman rho: -0.02). These results show the limitations of current state-of-the-art functional effect prediction models and underscore the challenges in modeling protein regions with distinct structural and mutational properties.

Overall, this generalization test as well as the plotting guidelines, the rBME metric and datasets from the previous sections provide a useful evaluation and could be helpful for the development of future models.

## 6 Conclusion

With our step-by-step model development approach building on and improving on previous methods, we develop a new functional effect predictor: ESM-Effect - an ESM2-fine-tuning architecture with an inductive bias regression head. ESM-Effect achieves SOTA performance across a range of DMSs by focusing on task-specific fine-tuning of PLM embeddings and not incorporating structure and MSA features.

Our model exhibits varying degrees of generalization when training and testing on distinct areas of the protein. This shows that further advancements are required before it may reliably inform treatments and deliver real-world benefits. We hope that our proposed benchmarking framework and the novel performance metric rBME may aid this progress by minimizing inflated results and facilitating robust comparisons between models.

Surprisingly, our analyses revealed that test performance remained almost constant across increasing model sizes during DMS fine-tuning and LoRA consistently matched the performance of full fine-tuning. These unexpected patterns diverge from typical natural (and protein) language model scaling behaviors and suggest that the PLM’s utility for DMS prediction may be fundamentally constrained by the biological depth of its pre-training approach. We hypothesize that only low-level, universal knowledge — largely invariant to model size — contributes meaningfully to DMS prediction. The performance plateau indicates that current pre-training paradigms struggle to capture the nuanced and detailed biological knowledge required for comprehensive functional effect prediction.

While current pre-training methods are effective at decoding sequence and structural aspects, they seem to fall short in capturing the complex biochemical reactions and interactions of proteins that are only weakly and implicitly encoded in sequence and structure. This suggests the need for new pre-training data sources and objectives (Li et al., 2024), capable of uncovering deeper biological insights.

Overall, our work highlights how efficient fine-tuning of PLMs achieves SOTA performance for functional effect prediction while being significantly less-compute intensive and sparing complex, multi-modal features. Our benchmarking framework emphasizes performance in rare, Gain-of-Function mutations with its relative Bin-Mean-Error metric while the generalization test provides a useful evaluation of transfer capabilities. We show as a proof-of-concept that ESM-Effect indeed exhibits generalization capabilities to some unseen regions which can be helpful for precision medicine and protein engineering applications.

## Author Contributions

**Moritz Glaser:** Conceptualization, Methodology, Software, Validation, Formal Analysis, Investigation, Data Curation, Writing - Original Draft, Visualization, Project Administration

**Johannes Brägelmann:** Supervision, Writing - Review & Editing, Conceptualization

## 7 Appendix

### 7.1 The Landscape of Pathogenicity Predictors

With the advent of artificial intelligence, advanced deep learning models (Krizhevsky et al., 2017) join the traditionally machine-learning-dominated landscape of mutation prediction models (Ioannidis et al., 2016; Adzhubei et al., 2010). The current landscape is characterized by two axes (cf. Figure 7):

**Figure 7:**
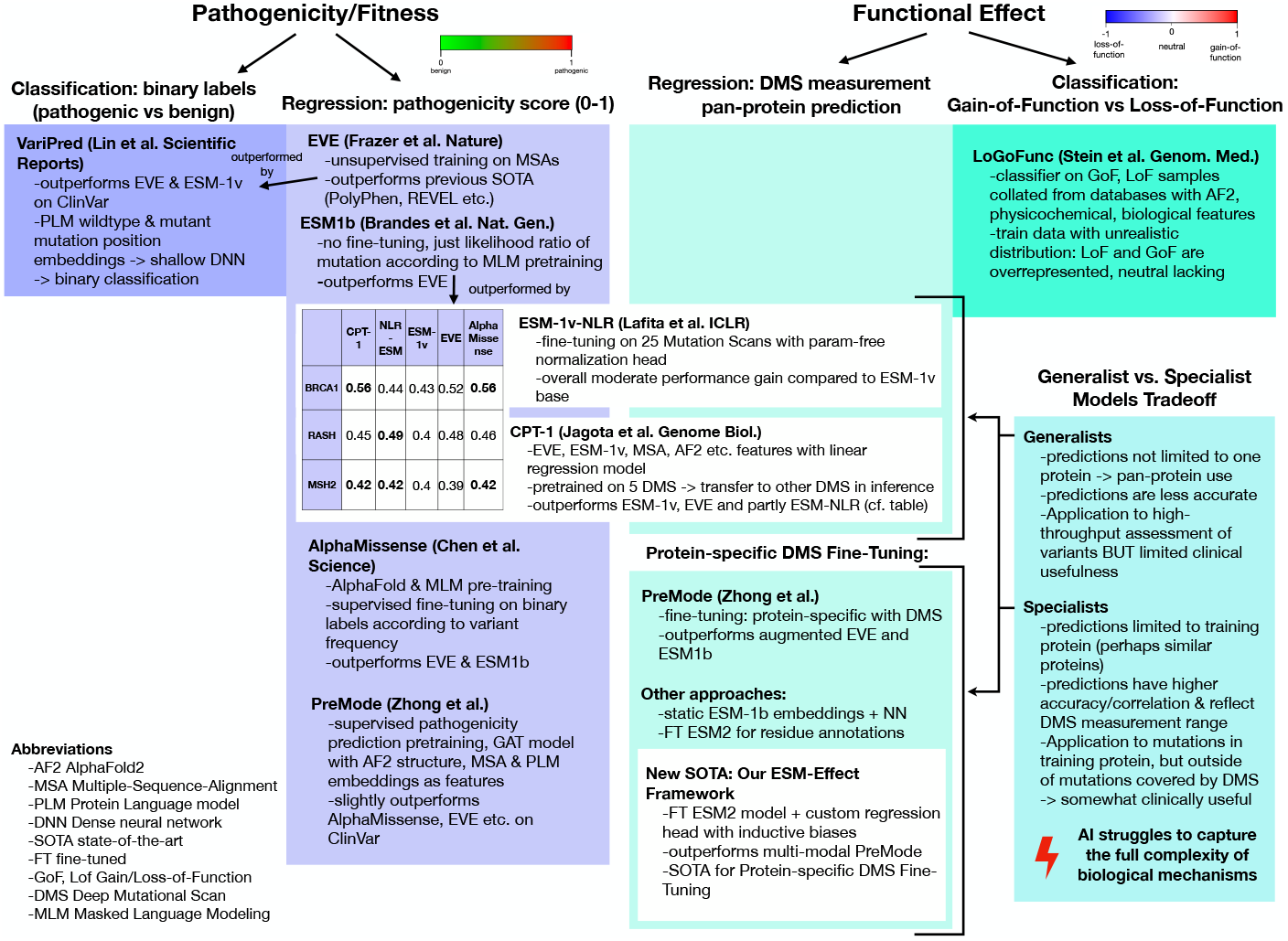
Survey of existing methods illustrating the trade-off between broadly applicable but less precise models and highly precise models limited to their training protein. Notably, the latter can produce high-quality predictions only for mutations within the same protein as the training DMS. Despite this limitation, such models remain valuable, as DMS datasets typically focus on specific protein domains and often contain incomplete data due to failed mutagenesis experiments.

- (a) whether the mutation effect is a unidirectional pathogenicity score or a bidirectional functional effect (i.e., increasing or decreasing a specific property or activity) and
- (b) whether the model performs classification or regression.

Most existing models focus on pathogenicity prediction (i.e., how physiological or wildtype-similar a mutation is) and use regression-based approaches. These models adopt a generalist strategy, scoring all possible variants across the (human) proteome (Cheng et al., 2023). However, pathogenicity predictors — whether trained on multiple DMS datasets, ClinVar annotations or physiological sequences — struggle to accurately predict the multi-faceted functional effects of specific mutations, such as rare gain-of-function enzyme mutations (Figure 8). This limitation arises from the biological complexity and specificity required for such tasks, which cannot be reliably captured by large-scale pre-training and the current architectures (Livesey & Marsh, 2023). However, clinical decision-making often depends on understanding the precise functional effect of mutations (i.e., increase/decrease of a specific protein property) (Iyer et al., 2023). Thus, functional effect predictors have been developed which are fine-tuned on DMS datasets achieving higher protein-specifity and accuracy yet are limited to unseen mutations and regions in their training protein. Current approaches rely on non-fine-tuned embeddings or intricate multi-modal features (Marquet et al., 2022; Zhong et al., 2024).

**Figure 8:**
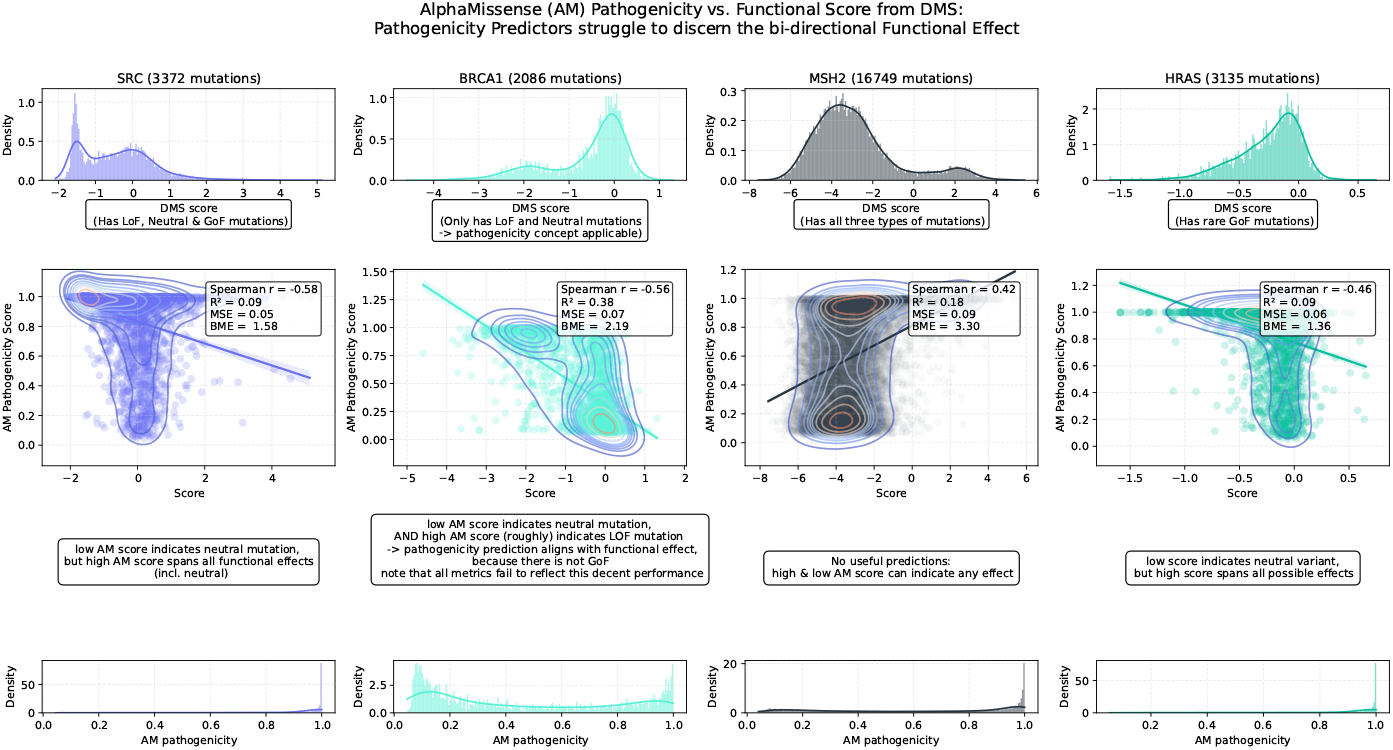
SOTA pathogenicity predictor AlphaMissense on DMS data. Note that the DMSs some-times not cover the entire protein sequence.

### 7.2 Protein modeling and pathogenicity prediction

Models like AlphaFold2 (AF2) predict protein structures from MSAs, capturing evolutionary information about residue interactions (Jumper et al., 2021) and Transformer-based Protein Language Models (PLMs), like ESM-1b and ESM2, learn protein semantics by predicting masked amino acids from evolutionary sequences (Rives et al., 2021; Lin et al., 2023; Rao et al., 2020). Because these models learn sequence and structure physiology they are directly applied to predict the lack thereof in form of the likelihood ratio of a mutant and wildtype residue (e.g., AlphaMissense, EVE building on MSAs (Cheng et al., 2023; Frazer et al., 2021) and pre-trained PLMs like ESM-1v (Meier et al., 2021; Brandes et al., 2023)). Some methods refine their training data with DMSs, which offer sufficient signal for pathogenicity despite heterogeneous properties across different DMSs. Examples include fine-tuning ESM-1v on 25 DMSs with a Normalized Log-odds Ratio (NLR) head (Lafita et al., 2024) and combining EVE, ESM-1v, and AF2 features in a regression model (Jagota et al., 2023). However, these methods struggle with multi-directional functional effects, particularly for Gain-of-Function mutations in DMSs like SNCA (Livesey & Marsh, 2023). In summary, while pathogenicity models effectively distinguish benign and pathogenic mutations, they fall short in predicting multi-dimensional functional effects as demonstrated in Figure 8.

### 7.3 Ablation and model analysis

#### Layer Probing

To investigate how the number of trainable layers affects performance, we retrained optimized ESM-Effect with a descending number of layers frozen: the results show that the number of frozen layers has no impact on test performance, as long as at least one layer remains unfrozen, allowing the model to adapt to the specific task (cf. Figure 9). Given that a single unfrozen layer can suffice for fine-tuning, we further explored whether its position within the network affects performance: the test performance remains consistent regardless of the unfrozen layer’s position. Even when only the first layer (immediately after the embedding layer) is unfrozen, it can still influence the subsequent layers, enabling the model to produce informative embeddings for the regression head at the final layer.

**Figure 9:**
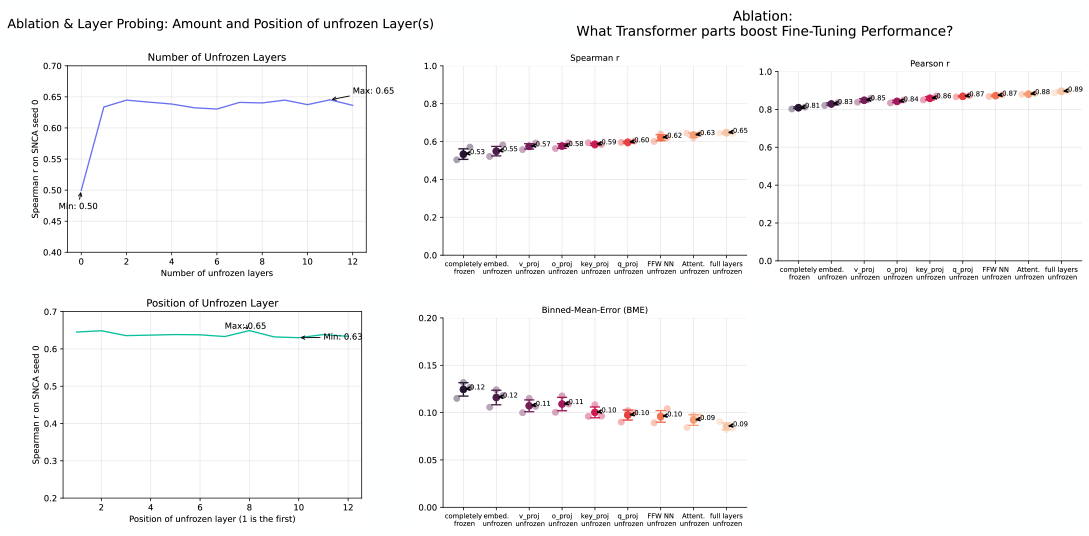
Ablation study of ESM-Effect: Fine-Tuning and Layer probing. Ablating Transformer parts of optimized ESM-Effect on 3 SNCA seeds.

#### Transformer Parts Ablation

To investigate which components of the Transformer architecture contribute most to performance, multiple models were trained with specific parts of the last two layers unfrozen. These include feed-forward layers, attention mechanisms, and individual components of the attention module (i.e., key, query, value and output projection layers). Performance (cf. Figure 9) increases progressively, starting from the embedding layer, followed by key, query, value and output projections, then the feed-forward and attention layers, and finally, the full last two layers.

This analysis suggests that ESM2 does not encode mutation-specific knowledge in individual layers, as it does for structural features such as contacts and binding sites (Vig et al., 2020). Fine-tuning performance is largely invariant to the position or number of fine-tuned layers, indicating that adaptation likely arises from task-specific tuning of the overall embeddings rather than mutation-specific mechanisms. Notably, the differences observed across Transformer components demonstrate the parameter efficiency of multi-head self-attention, which achieves competitive performance with approximately half the parameters of the feed-forward layers.

### 7.4 Experiments with Test-Time-Training (TTT)

As Bushuiev et al. (2024) showed, fine-tuning a pre-trained PLM backbone on a specific protein sequence that is used for a given inference task improves performance (Bushuiev et al., 2024). For instance, unsupervised mutation pathogenicity prediction from PLMs without a regression head benefited from TTT. Here, we sought to apply this technique to ESM-Effect using a similar approach for supevised functional effect prediction: first we customize (i.e., fine-tune) the ESM2 backbone on the protein sequence of the DMS. Then we train the backbone with the ESM-Effect head on top on a DMS. To customise the 35M ESM2 model, we the hyperparamter settings and training duration recommended by Bushuiev et al. (2024) for 35M ESM2. However, this led to rapid overfitting to the DMS sequence in this case: for the target DMS sequence and another (non-DMS protein family) sequence, we monitored the percentage of correctly predicted tokens and their probability when predicting each token in the sequence individually (with a mask for that token). We used this strategy to adjust the learning rate to maintain accuracy of the non-related sequence while achieving increased accuracy on the TTT/DMS sequence (cf. Figure 10). Based on the results we selected 1e-5 as optimal learning rate, customized the ESM2 backbone and trained ESM-Effect on three seeds of the SNCA DMS.

**Figure 10:**
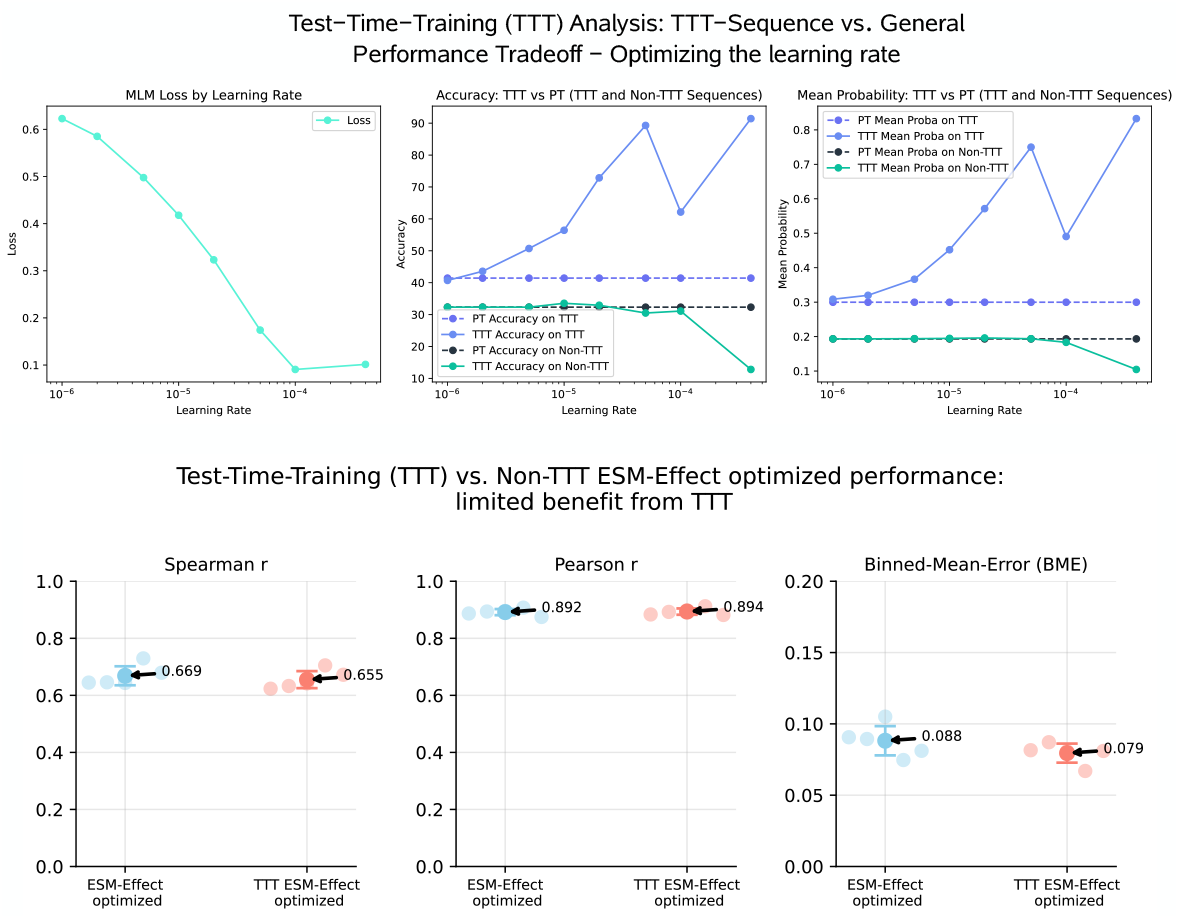
Top: Customizing ESM2 backbone on SNCA sequence while maintaining general knowledge to prevent catastrophic forgetting. Bottom: Mixed results from TTT fine-tuning on SNCA DMS.

Experiments with SNCA (seeds 0–2) reveal only minor performance differences between the non-TTT and TTT models, depending on the metric used. Consequently, no significant benefit from TTT is observed for the SNCA protein.

### 7.5 Generalization Test on distinct DMS

To examine whether ESM-Effect learns transferable features from one member of a protein family to another, we trained the model on the Glucokinase (GCK) DMS (with a 20% test split) and evaluated its performance on both the held-out GCK test set and an independent DMS from the SRC tyrosine kinase (Ahler et al., 2019).

Note that the distribution of the catalytic activity scores for both DMS is distinct, thus the i.i.d. (independent and identically distributed) assumption does not hold true (cf. Histogram Figure 11).

**Figure 11:**
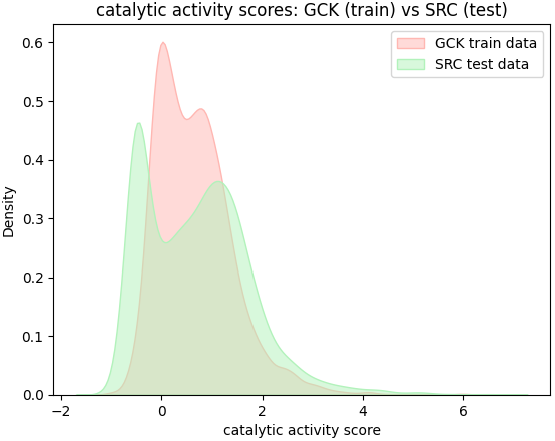
Catalytic score distributions for the GCK training and the SRC kinase testing DMS. The distinct distributions highlight the violation of the I.I.D. (independent and identically distributed) assumption. SRC DMS scores were min-max scaled to align with the range of GCK DMS values.

As illustrated in Figure 11, the catalytic activity score distributions for the GCK and SRC DMS datasets are distinct, violating the I.I.D. assumption fundamental to many machine learning models. This is biologically expected: despite both proteins being kinases, differences in their binding pockets and catalytic domains reflect their specialization for different substrates. Consequently, generalization from GCK to SRC is challenging.

While ESM-Effect achieves reasonable performance on the GCK test split (Spearman *ρ* = 0.67), it performs poorly on the SRC DMS (Spearman *ρ* = 0.03), underscoring the limited transferability of learned features across struc-turally and functionally divergent kinases.

### 7.6 Dissecting the Notion of a Wildtype Sequence

Over the course of ongoing evolution many different variants of sequences evolve and are selected for fitness. Thus, one fixed, unique “wild-type” sequence does not exist. Only different versions of sequences exist which have different properties. The term “mutation” and “variant” build on the arguable existence of one unique, static “wild-type” sequence in which one amino-acid is substituted forming the mutant sequence. Nonetheless, a physiological, natural sequence space exists comprising many functionally and fitness-regarding equivalent “wild-type” sequences which are curated in databases like UniProt (The UniProt Consortium et al., 2023), UniRef or SwissProt (Suzek et al., 2007; Boeckmann, 2003). These databases typically list one fixed, reference/”wild-type” sequence but also other isoforms. And different amino acid alterations in these physiological sequences may be viewed as mutations in contexts like precision medicine, where the wildtype sequence (space) for a given oncogene is established. In this light, the task of variant pathogenicity prediction equates to carving out the edges of the physiological sequence space. So the notion of one unique wild-type sequence is less applicable to variant pathogenicity prediction models, since the models learn a notion of physiological sequence spaces to which they compare a given sequence at inference. Yet they require a reference sequence (one version of the physiological wildtype) to compare the likelihood of the variant amino acid to: There is no effect without a reference to compare the effect to. The same logic applies to supervised, specialists models trained on DMSs. While we train models that only take the mutated sequence as input to predict the DMS score, the DMS score itself is being calculated by comparing the property of the cell expressing the mutant sequence to cells expressing the reference sequence. In general, variant prediction is not possible without a reference sequence (as part of the physiological sequence space).

### 7.7 Embedding Analysis

Seeking to understand how fine-tuned ESM2 embeddings compare to baseline ESM2 embeddings - the reason ESM-Effect outperforms PreMode - we obtained the embeddings for 100 GCK DMS test mutations from both models and analyzed them using the UMAP dimensionality reduction technique. However, there are no clusters and coloring the data points according to their catalytic activity does not show any relationship either. This might be due to the regression head’s importance in extracting meaningful features from the embedding (since it is trained with an order-of-magnitude higher learning rate).

### 7.8 Training Mean Embedding Models

Fine-tuning ESM2 models with a regression head using the mean sequence embedding presents unique challenges that do not arise when using the mutation position head. Notably, these issues are specific to the PTEN stability and enzyme activity DMS datasets and are not observed for SNCA or NUDT15. Training with the mean embedding often exhibits instability, characterized by spiking losses and abrupt fluctuations in performance. Additionally, convergence is slow, requiring more than 20–30 epochs, because the mean embedding condenses fine-grained information into a less informative representation, making it harder for the model to capture fine-grained mutation effects. Furthermore, the gradients from the regression head propagate less directly through the mean operation to the ESM2 model, compared to using the mutation position embedding, where the gradients flow directly from the head to the model parameters. This instability mainly applies to fine-tuning the full ESM2 model on the PTEN enzyme activity DMS compared to frozen or LoRA-based models. Therefore the PTEN enzyme activity comparison in Figure 2 is lacking given our limited compute resources in order to train enough models for enough epochs in the fully unfrozen setups.

### 7.9 Methods

#### 7.9.1 Training

A split the learning rate approach was used for ESM2 backbone and the regression head: ESM2 parameters are updated with a learning rate of 1e-5 and prediction head parameters by 1e-4 multiplied by the batch size (8), respectively. Dropout rate is set to 20% and we train for 10 epochs with a one cycle learning rate scheduler. AdamW was used with *β*_1_ = 0.9, *β*_2_ = 0.999, *ϵ* = 1e−8 and weight decay coefficient = 0.01. Training time for a DMS with 3k mutations and sequence length of 400 (e.g., PTEN) for 10 epochs is roughly 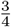 hour on a NVIDIA T4 GPU depending on evaluation and monitoring.

#### 7.9.2 Estimating Training Time and Resources of PreMode and ESM-Effect

While we were not able to run PreMode due to limited resources, the authors noted that the average DMS fine-tuning time for PreMode is 30 minutes on an NVIDIA A40 GPU which has 74.8 TFLOPS. ESM-Effect trains between 20 minutes and one hour on an NVIDIA T4 GPU with 8.1 TFLOPS. Correcting for the performance difference, PreMode trains for 4.5 T4 hours, while ESM-Effect trains for 0.67 hours (average). The ESM-Effect Speed Implementation caches the embeddings from the 10th layer in the first epoch and only performs forward-passes (and back-propagations) on the last two layers. Thus the average training time is about 5 minutes - an 8.4x speedup over ESM-Effect optimized and a 56.3x speedup over PreMode. Note that ESM-Effect can be further sped-up with FlashAttention (Dao et al., 2022) but requires NVIDIA Ampere GPUs or newer and does not work on Tesla GPUs.

While ESM-Effect only requires the model weights (ca. 200 MB), PreMode requires pre-computed ESM2 650M embeddings, AF2 structures and MSAs which amounts to roughly 60 gigabytes of compressed tar files. However, the memory requirements of ESM-Effect can increase during training depending on batch size and sequence length.

#### 7.9.3 Data

We used the same DMSs as in PreMode to compare performance: the same 20 percent test split with five (or three) different seeds was used for random splitting. Note that when using data from the PreMode repository the same .csv file contains scores for all properties of the DMS if there are multiple. As the score column names are not indicative of the measurement, and the same measurement type has different score column indices for different datasets we specify them here:

To evaluate generalization from training on GCK we use a DMS of the SRC kinase from MaveDB containing 3372 mutations (Ahler et al., 2019; Rubin et al., 2021). To adjust the scale of the score measurement from the SRC DMS to GCK DMS we use min-max scaling.

## References

Ivan A Adzhubei, Steffen Schmidt, Leonid Peshkin, Vasily E Ramensky, Anna Gerasimova, Peer Bork, Alexey S Kondrashov, and Shamil R Sunyaev. A method and server for predicting damaging missense mutations. Nature Methods, 7(4):248–249, April 2010. ISSN 1548-7091, 1548-7105. doi: 10.1038/nmeth0410-248. URL https://www.nature.com/articles/nmeth0410-248.

Ethan Ahler, Ames C. Register, Sujata Chakraborty, Linglan Fang, Emily M. Dieter, Katherine A. Sitko, Rama Subba Rao Vidadala, Bridget M. Trevillian, Martin Golkowski, Hannah Gelman, Jason J. Stephany, Alan F. Rubin, Ethan A. Merritt, Douglas M. Fowler, and Dustin J. Maly. A Combined Approach Reveals a Regulatory Mechanism Coupling Src’s Kinase Activity, Localization, and Phosphotransferase-Independent Functions. Molecular Cell, 74(2):393–408.e20, April 2019. ISSN 10972765. doi: 10.1016/j.molcel.2019.02.003. URL https://linkinghub.elsevier.com/retrieve/pii/S1097276519300930.

Joshua D. Backman, Alexander H. Li, Anthony Marcketta, Dylan Sun, Joelle Mbatchou, Michael D. Kessler, Christian Benner, Daren Liu, Adam E. Locke, Suganthi Balasubramanian, Ashish Yadav, Nilanjana Banerjee, Christopher E. Gillies, Amy Damask, Simon Liu, Xiaodong Bai, Alicia Hawes, Evan Maxwell, Lauren Gurski, Kyoko Watanabe, Jack A. Kosmicki, Veera Rajagopal, Jason Mighty, Regeneron Genetics Center, DiscovEHR, Marcus Jones, Lyndon Mitnaul, Eli Stahl, Giovanni Coppola, Eric Jorgenson, Lukas Habegger, William J. Salerno, Alan R. Shuldiner, Luca A. Lotta, John D. Overton, Michael N. Cantor, Jeffrey G. Reid, George Yancopoulos, Hyun M. Kang, Jonathan Marchini, Aris Baras, Gonçalo R. Abecasis, and Manuel A. R. Ferreira. Exome sequencing and analysis of 454,787 UK Biobank participants. Nature, 599(7886): 628–634, November 2021. ISSN 0028-0836, 1476-4687. doi: 10.1038/s41586-021-04103-z. URL https://www.nature.com/articles/s41586-021-04103-z.

Dan Biderman, Jacob Portes, Jose Javier Gonzalez Ortiz, Mansheej Paul, Philip Greengard, Connor Jennings, Daniel King, Sam Havens, Vitaliy Chiley, Jonathan Frankle, Cody Blakeney, and John P. Cunningham. LoRA Learns Less and Forgets Less, 2024. URL https://arxiv.org/abs/2405.09673. Version Number: 2.

B. Boeckmann. The SWISS-PROT protein knowledgebase and its supplement TrEMBL in 2003. Nucleic Acids Research, 31(1):365–370, January 2003. ISSN 13624962. doi: 10.1093/nar/gkg095. URL https://academic.oup.com/nar/article-lookup/doi/10.1093/nar/gkg095.

Nadav Brandes, Grant Goldman, Charlotte H. Wang, Chun Jimmie Ye, and Vasilis Ntranos. Genome-wide prediction of disease variant effects with a deep protein language model. Nature Genetics, 55(9):1512–1522, September 2023. ISSN 1061-4036, 1546-1718. doi: 10.1038/s41588-023-01465-0. URL https://www.nature.com/articles/s41588-023-01465-0.

Anton Bushuiev, Roman Bushuiev, Nikola Zadorozhny, Raman Samusevich, Hannes Stärk, Jiri Sedlar, Tomáš Pluskal, and Josef Sivic. Training on test proteins improves fitness, structure, and function prediction, 2024. URL https://arxiv.org/abs/2411.02109. Version Number: 1.

Jun Cheng, Guido Novati, Joshua Pan, Clare Bycroft, Akvilė Žemgulytė, Taylor Applebaum, Alexander Pritzel, Lai Hong Wong, Michal Zielinski, Tobias Sargeant, Rosalia G. Schneider, Andrew W. Senior, John Jumper, Demis Hassabis, Pushmeet Kohli, and Žiga Avsec. Accurate proteome-wide missense variant effect prediction with AlphaMissense. Science, 381(6664): eadg7492, September 2023. ISSN 0036-8075, 1095-9203. doi: 10.1126/science.adg7492. URL https://www.science.org/doi/10.1126/science.adg7492.

Xingyi Cheng, Bo Chen, Pan Li, Jing Gong, Jie Tang, and Le Song. Training compute-optimal protein language models. bioRxiv, 2024. doi: 10.1101/2024.06.06.597716. URL https://www.biorxiv.org/content/early/2024/06/09/2024.06.06.597716.

Tri Dao, Dan Fu, Stefano Ermon, Atri Rudra, and Christopher Ré. Flashattention: Fast and memory-efficient exact attention with io-awareness. In S. Koyejo, S. Mohamed, A. Agarwal, D. Belgrave, K. Cho, and A. Oh (eds.), Advances in Neural Information Processing Systems, volume 35, pp. 16344–16359. Curran Associates, Inc., 2022. URL https://proceedings.neurips.cc/paper_files/paper/2022/file/67d57c32e20fd0a7a302cb81d36e40d5-Paper-Conference.pdf.

Houssemeddine Derbel, Zhongming Zhao, and Qian Liu. Accurate prediction of functional effect of single amino acid variants with deep learning. Computational and Structural Biotechnology Journal, 21:5776–5784, 2023. ISSN 20010370. doi: 10.1016/j.csbj.2023.11.017. URL https://linkinghub.elsevier.com/retrieve/pii/S2001037023004312.

Alistair S Dunham and Pedro Beltrao. Exploring amino acid functions in a deep mutational landscape. Molecular Systems Biology, 17(7):e10305, July 2021. ISSN 1744-4292, 1744-4292. doi: 10.15252/msb.202110305. URL https://www.embopress.org/doi/10.15252/msb.202110305.

Daniel Esposito, Jochen Weile, Jay Shendure, Lea M. Starita, Anthony T. Papenfuss, Frederick P. Roth, Douglas M. Fowler, and Alan F. Rubin. MaveDB: an open-source platform to distribute and interpret data from multiplexed assays of variant effect. Genome Biology, 20(1):223, December 2019. ISSN 1474-760X. doi: 10.1186/s13059-019-1845-6. URL https://genomebiology.biomedcentral.com/articles/10.1186/s13059-019-1845-6.

Exome Aggregation Consortium, Monkol Lek, Konrad J. Karczewski, Eric V. Minikel, Kaitlin E. Samocha, Eric Banks, Timothy Fennell, Anne H. O’Donnell-Luria, James S. Ware, Andrew J. Hill, Beryl B. Cummings, Taru Tukiainen, Daniel P. Birnbaum, Jack A. Kosmicki, Laramie E. Duncan, Karol Estrada, Fengmei Zhao, James Zou, Emma Pierce-Hoffman, Joanne Berghout, David N. Cooper, Nicole Deflaux, Mark DePristo, Ron Do, Jason Flannick, Menachem Fromer, Laura Gauthier, Jackie Goldstein, Namrata Gupta, Daniel Howrigan, Adam Kiezun, Mitja I. Kurki, Ami Levy Moonshine, Pradeep Natarajan, Lorena Orozco, Gina M. Peloso, Ryan Poplin, Manuel A. Rivas, Valentin Ruano-Rubio, Samuel A. Rose, Douglas M. Ruderfer, Khalid Shakir, Peter D. Stenson, Christine Stevens, Brett P. Thomas, Grace Tiao, Maria T. Tusie-Luna, Ben Weisburd, Hong-Hee Won, Dongmei Yu, David M. Altshuler, Diego Ardissino, Michael Boehnke, John Danesh, Stacey Donnelly, Roberto Elosua, Jose C. Florez, Stacey B. Gabriel, Gad Getz, Stephen J. Glatt, Christina M. Hultman, Sekar Kathiresan, Markku Laakso, Steven McCarroll, Mark I. McCarthy, Dermot McGovern, Ruth McPherson, Benjamin M. Neale, Aarno Palotie, Shaun M. Purcell, Danish Saleheen, Jeremiah M. Scharf, Pamela Sklar, Patrick F. Sullivan, Jaakko Tuomilehto, Ming T. Tsuang, Hugh C. Watkins, James G. Wilson, Mark J. Daly, and Daniel G. MacArthur. Analysis of protein-coding genetic variation in 60,706 humans. Nature, 536(7616):285–291, August 2016. ISSN 0028-0836, 1476-4687. doi: 10.1038/nature19057. URL https://www.nature.com/articles/nature19057.

Jonathan Frazer, Pascal Notin, Mafalda Dias, Aidan Gomez, Joseph K. Min, Kelly Brock, Yarin Gal, and Debora S. Marks. Disease variant prediction with deep generative models of evolutionary data. Nature, 599(7883):91–95, November 2021. ISSN 0028-0836, 1476-4687. doi: 10.1038/s41586-021-04043-8. URL https://www.nature.com/articles/s41586-021-04043-8.

Hong Gao, Tobias Hamp, Jeffrey Ede, Joshua G. Schraiber, Jeremy McRae, Moriel Singer-Berk, Yanshen Yang, Anastasia S. D. Dietrich, Petko P. Fiziev, Lukas F. K. Kuderna, Laksshman Sundaram, Yibing Wu, Aashish Adhikari, Yair Field, Chen Chen, Serafim Batzoglou, Francois Aguet, Gabrielle Lemire, Rebecca Reimers, Daniel Balick, Mareike C. Janiak, Martin Kuhlwilm, Joseph D. Orkin, Shivakumara Manu, Alejandro Valenzuela, Juraj Bergman, Marjolaine Rousselle, Felipe Ennes Silva, Lidia Agueda, Julie Blanc, Marta Gut, Dorien De Vries, Ian Goodhead, R. Alan Harris, Muthuswamy Raveendran, Axel Jensen, Idriss S. Chuma, Julie E. Horvath, Christina Hvilsom, David Juan, Peter Frandsen, Fabiano R. De Melo, Fabrício Bertuol, Hazel Byrne, Iracilda Sampaio, Izeni Farias, João Valsecchi Do Amaral, Mariluce Messias, Maria N. F. Da Silva, Mihir Trivedi, Rogerio Rossi, Tomas Hrbek, Nicole Andriaholinirina, Clément J. Rabarivola, Alphonse Zaramody, Clifford J. Jolly, Jane Phillips-Conroy, Gregory Wilkerson, Christian Abee, Joe H. Simmons, Eduardo Fernandez-Duque, Sree Kanthaswamy, Fekadu Shiferaw, Dongdong Wu, Long Zhou, Yong Shao, Guojie Zhang, Julius D. Keyyu, Sascha Knauf, Minh D. Le, Esther Lizano, Stefan Merker, Arcadi Navarro, Thomas Bataillon, Tilo Nadler, Chiea Chuen Khor, Jessica Lee, Patrick Tan, Weng Khong Lim, Andrew C. Kitchener, Dietmar Zinner, Ivo Gut, Amanda Melin, Katerina Guschanski, Mikkel Heide Schierup, Robin M. D. Beck, Govindhaswamy Umapathy, Christian Roos, Jean P. Boubli, Monkol Lek, Shamil Sunyaev, Anne O’Donnell-Luria, Heidi L. Rehm, Jinbo Xu, Jeffrey Rogers, Tomas Marques-Bonet, and Kyle Kai-How Farh. The landscape of tolerated genetic variation in humans and primates. Science, 380(6648):eabn8153, June 2023. ISSN 0036-8075, 1095-9203. doi: 10.1126/science.abn8197. URL https://www.science.org/doi/10.1126/science.abn8197.

Nilah M. Ioannidis, Joseph H. Rothstein, Vikas Pejaver, Sumit Middha, Shannon K. McDonnell, Saurabh Baheti, Anthony Musolf, Qing Li, Emily Holzinger, Danielle Karyadi, Lisa A. Cannon-Albright, Craig C. Teerlink, Janet L. Stanford, William B. Isaacs, Jianfeng Xu, Kathleen A. Cooney, Ethan M. Lange, Johanna Schleutker, John D. Carpten, Isaac J. Powell, Olivier Cussenot, Geraldine Cancel-Tassin, Graham G. Giles, Robert J. MacInnis, Christiane Maier, Chih-Lin Hsieh, Fredrik Wiklund, William J. Catalona, William D. Foulkes, Diptasri Mandal, Rosalind A. Eeles, Zsofia Kote-Jarai, Carlos D. Bustamante, Daniel J. Schaid, Trevor Hastie, Elaine A. Ostrander, Joan E. Bailey-Wilson, Predrag Radivojac, Stephen N. Thibodeau, Alice S. Whittemore, and Weiva Sieh. REVEL: An Ensemble Method for Predicting the Pathogenicity of Rare Missense Variants. The American Journal of Human Genetics, 99(4):877–885, October 2016. ISSN 00029297. doi: 10.1016/j.ajhg.2016.08.016. URL https://linkinghub.elsevier.com/retrieve/pii/S0002929716303706.

Sudarshan R Iyer, Kevin Nusser, Kristen Jones, Pushkar Shinde, Clare Keddy, Catherine Z Beach, Erin Aguero, Jeremy Force, Ujwal Shinde, and Monika A Davare. Discovery of oncogenic ROS1 missense mutations with sensitivity to tyrosine kinase inhibitors. EMBO Molecular Medicine, 15(10):e17367, October 2023. ISSN 1757-4676, 1757-4684. doi: 10.15252/emmm.202217367. URL https://www.embopress.org/doi/10.15252/emmm.202217367.

Milind Jagota, Chengzhong Ye, Carlos Albors, Ruchir Rastogi, Antoine Koehl, Nilah Ioannidis, and Yun S. Song. Cross-protein transfer learning substantially improves disease variant prediction. Genome Biology, 24(1):182, August 2023. ISSN 1474-760X. doi: 10.1186/s13059-023-03024-6. URL https://genomebiology.biomedcentral.com/articles/10.1186/s13059-023-03024-6.

John Jumper, Richard Evans, Alexander Pritzel, Tim Green, Michael Figurnov, Olaf Ronneberger, Kathryn Tunyasuvunakool, Russ Bates, Augustin Žídek, Anna Potapenko, Alex Bridgland, Clemens Meyer, Simon A. A. Kohl, Andrew J. Ballard, Andrew Cowie, Bernardino Romera-Paredes, Stanislav Nikolov, Rishub Jain, Jonas Adler, Trevor Back, Stig Petersen, David Reiman, Ellen Clancy, Michal Zielinski, Martin Steinegger, Michalina Pacholska, Tamas Berghammer, Sebastian Bodenstein, David Silver, Oriol Vinyals, Andrew W. Senior, Koray Kavukcuoglu, Pushmeet Kohli, and Demis Hassabis. Highly accurate protein structure prediction with AlphaFold. Nature, 596(7873):583–589, August 2021. ISSN 0028-0836, 1476-4687. doi: 10.1038/s41586-021-03819-2. URL https://www.nature.com/articles/s41586-021-03819-2.

Jared Kaplan, Sam McCandlish, Tom Henighan, Tom B. Brown, Benjamin Chess, Rewon Child, Scott Gray, Alec Radford, Jeffrey Wu, and Dario Amodei. Scaling Laws for Neural Language Models, 2020. URL https://arxiv.org/abs/2001.08361. Version Number: 1.

Konrad J. Karczewski, Laurent C. Francioli, Grace Tiao, Beryl B. Cummings, Jessica Alföldi, Qingbo Wang, Ryan L. Collins, Kristen M. Laricchia, Andrea Ganna, Daniel P. Birnbaum, Laura D. Gauthier, Harrison Brand, Matthew Solomonson, Nicholas A. Watts, Daniel Rhodes, Moriel Singer-Berk, Eleina M. England, Eleanor G. Seaby, Jack A. Kosmicki, Raymond K. Walters, Katherine Tashman, Yossi Farjoun, Eric Banks, Timothy Poterba, Arcturus Wang, Cotton Seed, Nicola Whiffin, Jessica X. Chong, Kaitlin E. Samocha, Emma Pierce-Hoffman, Zachary Zappala, Anne H. O’Donnell-Luria, Eric Vallabh Minikel, Ben Weisburd, Monkol Lek, James S. Ware, Christopher Vittal, Irina M. Armean, Louis Bergelson, Kristian Cibulskis, Kristen M. Connolly, Miguel Covarrubias, Stacey Donnelly, Steven Ferriera, Stacey Gabriel, Jeff Gentry, Namrata Gupta, Thibault Jeandet, Diane Kaplan, Christopher Llanwarne, Ruchi Munshi, Sam Novod, Nikelle Petrillo, David Roazen, Valentin Ruano-Rubio, Andrea Saltzman, Molly Schleicher, Jose Soto, Kathleen Tibbetts, Charlotte Tolonen, Gordon Wade, Michael E. Talkowski, Genome Aggregation Database Consortium, Carlos A. Aguilar Salinas, Tariq Ahmad, Christine M. Albert, Diego Ardissino, Gil Atzmon, John Barnard, Laurent Beaugerie, Emelia J. Benjamin, Michael Boehnke, Lori L. Bonnycastle, Erwin P. Bottinger, Donald W. Bowden, Matthew J. Bown, John C. Chambers, Juliana C. Chan, Daniel Chasman, Judy Cho, Mina K. Chung, Bruce Cohen, Adolfo Correa, Dana Dabelea, Mark J. Daly, Dawood Darbar, Ravindranath Duggirala, Joseé Dupuis, Patrick T. Ellinor, Roberto Elosua, Jeanette Erdmann, Tonu Esko, Martti Färkkilä, Jose Florez, Andre Franke, Gad Getz, Benjamin Glaser, Stephen J. Glatt, David Goldstein, Clicerio Gonzalez, Leif Groop, Christopher Haiman, Craig Hanis, Matthew Harms, Mikko Hiltunen, Matti M. Holi, Christina M. Hultman, Mikko Kallela, Jaakko Kaprio, Sekar Kathiresan, Bong-Jo Kim, Young Jin Kim, George Kirov, Jaspal Kooner, Seppo Koskinen, Harlan M. Krumholz, Subra Kugathasan, Soo Heon Kwak, Markku Laakso, Terho Lehtimäki, Ruth J. F. Loos, Steven A. Lubitz, Ronald C. W. Ma, Daniel G. MacArthur, Jaume Marrugat, Kari M. Mattila, Steven McCarroll, Mark I. McCarthy, Dermot McGovern, Ruth McPherson, James B. Meigs, Olle Melander, Andres Metspalu, Benjamin M. Neale, Peter M. Nilsson, Michael C. O’Donovan, Dost Ongur, Lorena Orozco, Michael J. Owen, Colin N. A. Palmer, Aarno Palotie, Kyong Soo Park, Carlos Pato, Ann E. Pulver, Nazneen Rahman, Anne M. Remes, John D. Rioux, Samuli Ripatti, Dan M. Roden, Danish Saleheen, Veikko Salomaa, Nilesh J. Samani, Jeremiah Scharf, Heribert Schunkert, Moore B. Shoemaker, Pamela Sklar, Hilkka Soininen, Harry Sokol, Tim Spector, Patrick F. Sullivan, Jaana Suvisaari, E. Shyong Tai, Yik Ying Teo, Tuomi Tiinamaija, Ming Tsuang, Dan Turner, Teresa Tusie-Luna, Erkki Vartiainen, Marquis P. Vawter, James S. Ware, Hugh Watkins, Rinse K. Weersma, Maija Wessman, James G. Wilson, Ramnik J. Xavier, Benjamin M. Neale, Mark J. Daly, and Daniel G. MacArthur. The mutational constraint spectrum quantified from variation in 141,456 humans. Nature, 581(7809): 434–443, May 2020. ISSN 0028-0836, 1476-4687. doi: 10.1038/s41586-020-2308-7. URL https://www.nature.com/articles/s41586-020-2308-7.

Alex Krizhevsky, Ilya Sutskever, and Geoffrey E. Hinton. ImageNet classification with deep convolutional neural networks. Communications of the ACM, 60(6):84–90, May 2017. ISSN 0001-0782, 1557-7317. doi: 10.1145/3065386. URL https://dl.acm.org/doi/10.1145/3065386.

Dimitri M Kullmann and Michael G Hanna. Neurological disorders caused by inherited ion-channel mutations. The Lancet Neurology, 1(3):157–166, July 2002. ISSN 14744422. doi: 10.1016/S1474-4422(02)00071-6. URL https://linkinghub.elsevier.com/retrieve/pii/S1474442202000716.

Aleix Lafita, Ferran Gonzalez, Mahmoud Hossam, Paul Smyth, Jacob Deasy, Ari Allyn-Feuer, Daniel Seaton, and Stephen Young. Fine-tuning Protein Language Models with Deep Mutational Scanning improves Variant Effect Prediction, 2024. URL https://arxiv.org/abs/2405.06729. Version Number: 1.

Melissa J Landrum, Jennifer M Lee, Mark Benson, Garth R Brown, Chen Chao, Shanmuga Chitipiralla, Baoshan Gu, Jennifer Hart, Douglas Hoffman, Wonhee Jang, Karen Karapetyan, Kenneth Katz, Chunlei Liu, Zenith Maddipatla, Adriana Malheiro, Kurt McDaniel, Michael Ovetsky, George Riley, George Zhou, J Bradley Holmes, Brandi L Kattman, and Donna R Maglott. ClinVar: improving access to variant interpretations and supporting evidence. Nucleic Acids Research, 46(D1):D1062–D1067, January 2018. ISSN 0305-1048, 1362-4962. doi: 10.1093/nar/gkx1153. URL http://academic.oup.com/nar/article/46/D1/D1062/4641904.

Jonas Leichsenring, Peter Horak, Simon Kreutzfeldt, Christoph Heining, Petros Christopoulos, Anna-Lena Volckmar, Olaf Neumann, Martina Kirchner, Carolin Ploeger, Jan Budczies, Christoph E. Heilig, Barbara Hutter, Martina Fröhlich, Sebastian Uhrig, Daniel Kazdal, Michael Allgäuer, Alexander Harms, Eugen Rempel, Ulrich Lehmann, Michael Thomas, Nicole Pfarr, Ninel Azoitei, Irina Bonzheim, Ralf Marienfeld, Peter Möller, Martin Werner, Falko Fend, Melanie Boerries, Nikolas Von Bubnoff, Silke Lassmann, Thomas Longerich, Michael Bitzer, Thomas Seufferlein, Nisar Malek, Wilko Weichert, Peter Schirmacher, Roland Penzel, Volker Endris, Benedikt Brors, Frederick Klauschen, Hanno Glimm, Stefan Fröhling, and Albrecht Stenzinger. Variant classification in precision oncology. International Journal of Cancer, 145 (11):2996–3010, December 2019. ISSN 0020-7136, 1097-0215. doi: 10.1002/ijc.32358. URL https://onlinelibrary.wiley.com/doi/10.1002/ijc.32358.

Francesca-Zhoufan Li, Ava P. Amini, Yisong Yue, Kevin K. Yang, and Alex X. Lu. Feature Reuse and Scaling: Understanding Transfer Learning with Protein Language Models, February 2024. URL http://biorxiv.org/lookup/doi/10.1101/2024.02.05.578959.

Zeming Lin, Halil Akin, Roshan Rao, Brian Hie, Zhongkai Zhu, Wenting Lu, Nikita Smetanin, Robert Verkuil, Ori Kabeli, Yaniv Shmueli, Allan Dos Santos Costa, Maryam Fazel-Zarandi, Tom Sercu, Salvatore Candido, and Alexander Rives. Evolutionary-scale prediction of atomic-level protein structure with a language model. Science, 379(6637):1123–1130, March 2023. ISSN 0036-8075, 1095-9203. doi: 10.1126/science.ade2574. URL https://www.science.org/doi/10.1126/science.ade2574.

Benjamin J Livesey and Joseph A Marsh. Updated benchmarking of variant effect predictors using deep mutational scanning. Molecular Systems Biology, 19(8):e11474, August 2023. ISSN 1744-4292, 1744-4292. doi: 10.15252/msb.202211474. URL https://www.embopress.org/doi/10.15252/msb.202211474.

Céline Marquet, Michael Heinzinger, Tobias Olenyi, Christian Dallago, Kyra Erckert, Michael Bernhofer, Dmitrii Nechaev, and Burkhard Rost. Embeddings from protein language models predict conservation and variant effects. Human Genetics, 141(10):1629–1647, October 2022. ISSN 0340-6717, 1432-1203. doi: 10.1007/s00439-021-02411-y. URL https://link.springer.com/10.1007/s00439-021-02411-y.

Joaquin Mateo, Lotte Steuten, Philippe Aftimos, Fabrice André, Mark Davies, Elena Garralda, Jan Geissler, Don Husereau, Iciar Martinez-Lopez, Nicola Normanno, Jorge S. Reis-Filho, Stephen Stefani, David M. Thomas, C. Benedikt Westphalen, and Emile Voest. Delivering precision oncology to patients with cancer. Nature Medicine, 28(4):658–665, April 2022. ISSN 1078-8956, 1546-170X. doi: 10.1038/s41591-022-01717-2. URL https://www.nature.com/articles/s41591-022-01717-2.

Joshua Meier, Roshan Rao, Robert Verkuil, Jason Liu, Tom Sercu, and Alexander Rives. Language models enable zero-shot prediction of the effects of mutations on protein function, July 2021. URL http://biorxiv.org/lookup/doi/10.1101/2021.07.09.450648.

Marco Necci, Damiano Piovesan, Zsuzsanna Dosztányi, and Silvio C.E Tosatto. MobiDB-lite: fast and highly specific consensus prediction of intrinsic disorder in proteins. Bioinformatics, 33(9):1402–1404, May 2017. ISSN 1367-4803, 1367-4811. doi: 10.1093/bioinformatics/btx015. URL https://academic.oup.com/bioinformatics/article/33/9/1402/2908909.

Pascal Notin, Aaron W. Kollasch, Daniel Ritter, Lood Van Niekerk, Steffanie Paul, Hansen Spinner, Nathan Rollins, Ada Shaw, Ruben Weitzman, Jonathan Frazer, Mafalda Dias, Dinko Franceschi, Rose Orenbuch, Yarin Gal, and Debora S. Marks. ProteinGym: Large-Scale Benchmarks for Protein Design and Fitness Prediction, December 2023. URL http://biorxiv.org/lookup/doi/10.1101/2023.12.07.570727.

Clémence T. B. Pasmans, Bastiaan B. J. Tops, Elisabeth M. P. Steeghs, Veerle M. H. Coupé, Katrien Grünberg, Eiko K De Jong, Ed M. D. Schuuring, Stefan M. Willems, Marjolijn J. L. Ligtenberg, Valesca P. Retel, Hans Van Snellenberg, Ewart De Bruijn, Edwin Cuppen, and Geert W. J. Frederix. Micro-costing diagnostics in oncology: from single-gene testing to whole-genome sequencing. Expert Review of Pharmacoeconomics & Outcomes Research, 21(3):413–414, May 2021. ISSN 1473-7167, 1744-8379. doi: 10.1080/14737167.2021.1917385. URL https://www.tandfonline.com/doi/full/10.1080/14737167.2021.1917385.

Roshan Rao, Joshua Meier, Tom Sercu, Sergey Ovchinnikov, and Alexander Rives. Transformer protein language models are unsupervised structure learners, December 2020. URL http://biorxiv.org/lookup/doi/10.1101/2020.12.15.422761.

Alexander Rives, Joshua Meier, Tom Sercu, Siddharth Goyal, Zeming Lin, Jason Liu, Demi Guo, Myle Ott, C. Lawrence Zitnick, Jerry Ma, and Rob Fergus. Biological structure and function emerge from scaling unsupervised learning to 250 million protein sequences. Proceedings of the National Academy of Sciences, 118(15):e2016239118, April 2021. ISSN 0027-8424, 1091-6490. doi: 10.1073/pnas.2016239118. URL https://pnas.org/doi/full/10.1073/pnas.2016239118.

Alan F Rubin, Joseph K Min, Nathan J Rollins, Estelle Y Da, Daniel Esposito, Matthew Harrington, Jeremy Stone, Aisha Haley Bianchi, Mafalda Dias, Jonathan Frazer, Yunfan Fu, Molly Gallaher, Iris Li, Olivia Moscatelli, Jesslyn Yl Ong, Joshua E Rollins, Matthew J Wakefield, Shenyi Sunny Ye, Amy Tam, Abbye E McEwen, Lea M Starita, Vanessa L Bryant, Debora S Marks, and Douglas M Fowler. MaveDB v2: a curated community database with over three million variant effects from multiplexed functional assays, November 2021. URL http://biorxiv.org/lookup/doi/10.1101/2021.11.29.470445.

Ali Saadat and Jacques Fellay. Fine-tuning the ESM2 protein language model to understand the functional impact of missense variants, 2024. URL https://arxiv.org/abs/2410.10919. Version Number: 1.

Robert Schmirler, Michael Heinzinger, and Burkhard Rost. Fine-tuning protein language models boosts predictions across diverse tasks. Nature Communications, 15(1):7407, August 2024. ISSN 2041-1723. doi: 10.1038/s41467-024-51844-2. URL https://www.nature.com/articles/s41467-024-51844-2.

Amelie Schreiber. Esmbind and qbind: Lora, qlora, and esm-2 for predicting binding sites and post translational modification. bioRxiv, 2023. doi: 10.1101/2023.11.13.566930. URL https://www.biorxiv.org/content/early/2023/11/14/2023.11.13.566930.

David Stein, Meltem Ece Kars, Yiming Wu, Çigdem Sevim Bayrak, Peter D. Stenson, David N. Cooper, Avner Schlessinger, and Yuval Itan. Genome-wide prediction of pathogenic gain- and loss-of-function variants from ensemble learning of a diverse feature set. Genome Medicine, 15(1):103, November 2023. ISSN 1756-994X. doi: 10.1186/s13073-023-01261-9. URL https://genomemedicine.biomedcentral.com/articles/10.1186/s13073-023-01261-9.

Baris E. Suzek, Hongzhan Huang, Peter McGarvey, Raja Mazumder, and Cathy H. Wu. UniRef: comprehensive and non-redundant UniProt reference clusters. Bioinformatics, 23 (10):1282–1288, May 2007. ISSN 1367-4811, 1367-4803. doi: 10.1093/bioinformatics/btm098. URL https://academic.oup.com/bioinformatics/article/23/10/1282/197795.

The UniProt Consortium, Alex Bateman, Maria-Jesus Martin, Sandra Orchard, Michele Magrane, Shadab Ahmad, Emanuele Alpi, Emily H Bowler-Barnett, Ramona Britto, Hema Bye-A-Jee, Austra Cukura, Paul Denny, Tunca Dogan, ThankGod Ebenezer, Jun Fan, Penelope Garmiri, Leonardo Jose Da Costa Gonzales, Emma Hatton-Ellis, Abdulrahman Hussein, Alexandr Ignatchenko, Giuseppe Insana, Rizwan Ishtiaq, Vishal Joshi, Dushyanth Jyothi, Swaathi Kandasaamy, Antonia Lock, Aurelien Luciani, Marija Lugaric, Jie Luo, Yvonne Lussi, Alistair MacDougall, Fabio Madeira, Mahdi Mahmoudy, Alok Mishra, Katie Moulang, Andrew Nightingale, Sangya Pundir, Guoying Qi, Shriya Raj, Pedro Raposo, Daniel L Rice, Rabie Saidi, Rafael Santos, Elena Speretta, James Stephenson, Prabhat Totoo, Edward Turner, Nidhi Tyagi, Preethi Vasudev, Kate Warner, Xavier Watkins, Rossana Zaru, Hermann Zellner, Alan J Bridge, Lucila Aimo, Ghislaine Argoud-Puy, Andrea H Auchincloss, Kristian B Axelsen, Parit Bansal, Delphine Baratin, Teresa M Batista Neto, Marie-Claude Blatter, Jerven T Bolleman, Emmanuel Boutet, Lionel Breuza, Blanca Cabrera Gil, Cristina Casals-Casas, Kamal Chikh Echioukh, Elisabeth Coudert, Beatrice Cuche, Edouard De Castro, Anne Estreicher, Maria L Famiglietti, Marc Feuermann, Elisabeth Gasteiger, Pascale Gaudet, Sebastien Gehant, Vivienne Gerritsen, Arnaud Gos, Nadine Gruaz, Chantal Hulo, Nevila Hyka-Nouspikel, Florence Jungo, Arnaud Kerhornou, Philippe Le Mercier, Damien Lieberherr, Patrick Masson, Anne Morgat, Venkatesh Muthukrishnan, Salvo Paesano, Ivo Pedruzzi, Sandrine Pilbout, Lucille Pourcel, Sylvain Poux, Monica Pozzato, Manuela Pruess, Nicole Redaschi, Catherine Rivoire, Christian J A Sigrist, Karin Sonesson, Shyamala Sundaram, Cathy H Wu, Cecilia N Arighi, Leslie Arminski, Chuming Chen, Yongxing Chen, Hongzhan Huang, Kati Laiho, Peter McGarvey, Darren A Natale, Karen Ross, C R Vinayaka, Qinghua Wang, Yuqi Wang, and Jian Zhang. UniProt: the Universal Protein Knowledgebase in 2023. Nucleic Acids Research, 51(D1):D523–D531, January 2023. ISSN 0305-1048, 1362-4962. doi: 10.1093/nar/gkac1052. URL https://academic.oup.com/nar/article/51/D1/D523/6835362.

Jesse Vig, Ali Madani, Lav R. Varshney, Caiming Xiong, Richard Socher, and Nazneen Fatema Rajani. BERTology Meets Biology: Interpreting Attention in Protein Language Models, 2020. URL https://arxiv.org/abs/2006.15222. Version Number: 3.

Guojie Zhong and Yufeng Shen. Representation of missense variants for predicting modes of action. Machine Learning for Structural Biology Workshop, NeurIPS, 2022.

Guojie Zhong, Yige Zhao, Demi Zhuang, Wendy K Chung, and Yufeng Shen. PreMode predicts mode-of-action of missense variants by deep graph representation learning of protein sequence and structural context, February 2024. URL http://biorxiv.org/lookup/doi/10.1101/2024.02.20.581321.

